# Touch sensation requires the mechanically-gated ion channel Elkin1

**DOI:** 10.1101/2023.06.09.544247

**Authors:** Sampurna Chakrabarti, Jasmin D. Klich, Mohammed A. Khallaf, Oscar Sánchez-Carranza, Zuzanna M. Baran, Alice Rossi, Angela Tzu-Lun Huang, Tobias Pohl, Raluca Fleischer, Carina Fürst, Annette Hammes, Valérie Bégay, Hanna Hörnberg, Kate Poole, Gary R. Lewin

## Abstract

The slightest touch to the skin initiates tactile perception that is almost immediate^1^. The extraordinary speed of touch perception is enabled by mechanically-activated ion channels, the opening of which excites the endings of sensory neurons innervating the skin to initiate sensation. Here we identify a new mechanically-activated ion channel, Elkin1^2^, that, when ablated in mice, leads to a profound behavioural touch insensitivity. Touch insensitivity in *Elkin1*^*-/-*^ mice was caused by a loss of mechanically-activated currents (MA-currents) in around half of all sensory neurons that are activated by light touch (low threshold mechanoreceptors, LTMRs). Reintroduction of Elkin1 into sensory neurons from *Elkin1*^*-/-*^ mice acutely restored MA-currents. Piezo2^3–6^ is an established mechanosensitive ion channel required for touch sensation. In mice genetic ablation of *Piezo2* renders many, but not all, LTMRs insensitive to mechanical force^4,5,7^. Here we show that Elkin1 underpins PIEZO2-independent touch sensation. Additionally, we find that Elkin1 is present in many nociceptive sensory neurons which detect potentially damaging and painful mechanical force. These nociceptors depend on *Elkin1* for effectively communicating information on sustained noxious mechanical forces. We further identified molecular and functional interactions between the known mechanotransduction protein Stoml3^8,9^ and Elkin1 ion channels. Our data identify Elkin1 as a novel core component of touch transduction in mammals. The specific sensory deficits exhibited by *Elkin1*^*-/-*^ mice make Elkin1 a highly desirable target that could be harnessed to treat somatic sensory disorders including pain.

## Main text

Touch sensation is fundamental to our sense of self, social interactions and exploration of the tactile world^1,10^. Sensation is initiated at specialized end-organs in the skin, innervated by low threshold mechanoreceptors (LTMRs) with their cell bodies in the dorsal root ganglion (DRG). The peripheral endings of LTMRs are equipped with mechanically-gated ion channels that can be opened by tiny forces to initiate and enable touch perception^11,12^. The mechanically-gated ion channel Piezo2 is expressed by most sensory neurons in the DRG^3^ and in its absence, around half of LTMRs no longer respond to mechanical stimuli^4,5,7^. The DRG also contains so-called nociceptors, sensory neurons specialized to detect potentially damaging and painful stimuli, including intense mechanical force^11^. Many nociceptors express Piezo2 channels, but remain mechanosensitive in its absence^5^. The preservation of largely normal nociceptor mechanosensitivity in the absence of Piezo2 channels led us to search for other mechanically-gated ion channels that could account for Piezo2-independent sensory mechanotransduction.

We previously identified ELKIN1 (TMEM87A) as a protein that is both necessary and sufficient to confer mechanosensitivity to highly metastatic human melanoma cells^2^. Cryo-EM structures of human Elkin1 recently revealed a monomeric 7 transmembrane protein with an N-terminal extracellular Golgi-dynamics domain fold (GOLD-domain)^13^. A second higher resolution structure recently identified a cation conduction pathway through the protein^14^. We overexpressed Elkin1 in HEK293T cells lacking Piezo1 channels (HEK293T^*Piezo1-/-*^ cells)^15^ and found large poking-induced mechanically-activated currents (MA-currents) in a majority of transfected cells (Fig. 1a,b). Cells were plated on laminin 511 and poly-L-lysine (PLL), a substrate that supports increased mechanosensitivity^16^; untransfected cells showed no poking-induced currents. Elkin1-dependent currents were rapidly-adapting (RA) with fast inactivation time constants (<10 ms) similar to those of Piezo2 ion channels^3,17^ (Fig. 1 a,b,c). Using substrate deflection via pillar arrays^2,17^, we also found robust mechanically-activated currents in HEK293T^*Piezo1-/-*^ cells, transfected with *Elkin1* expression constructs, but also in cells transfected with *Elkin1* lacking the N-terminal GOLD-domain (*Elk1Δ170*) (Fig. 1c,d Extended data Fig. 1a)^2^. Most of the pillar evoked currents were rapidly-adapting (RA, inactivation <10 ms) or intermediately adapting (IA, inactivation between 10-50 ms). Measurements of pillar gated currents at different holding potentials revealed a linear current-voltage relation with a reversal potential of 0 mV for both *Elkin1* and *Elk1Δ170* transfected cells (Fig. 1e). Therefore, the GOLD-domain is not necessary for mechanical activation of Elkin1. Elkin1 currents showed a distinctive pharmacological profile, being highly sensitive to the non-specific pore-blocker Gd^3+^, but barely affected by concentrations of ruthenium red (30 μM) that completely block other mechanosensitive channels like Piezo1 and Piezo2^3^ (Fig. 1f, Extended data Fig 1b). Additionally, in agreement with a recent report^14^ we found that cells expressing mouse Elkin1 display prominent leak currents at very positive (> +60 mV) and very negative potentials (< -100 mV) (Fig 1g, Extended data Fig 1c).

**Figure 1:**
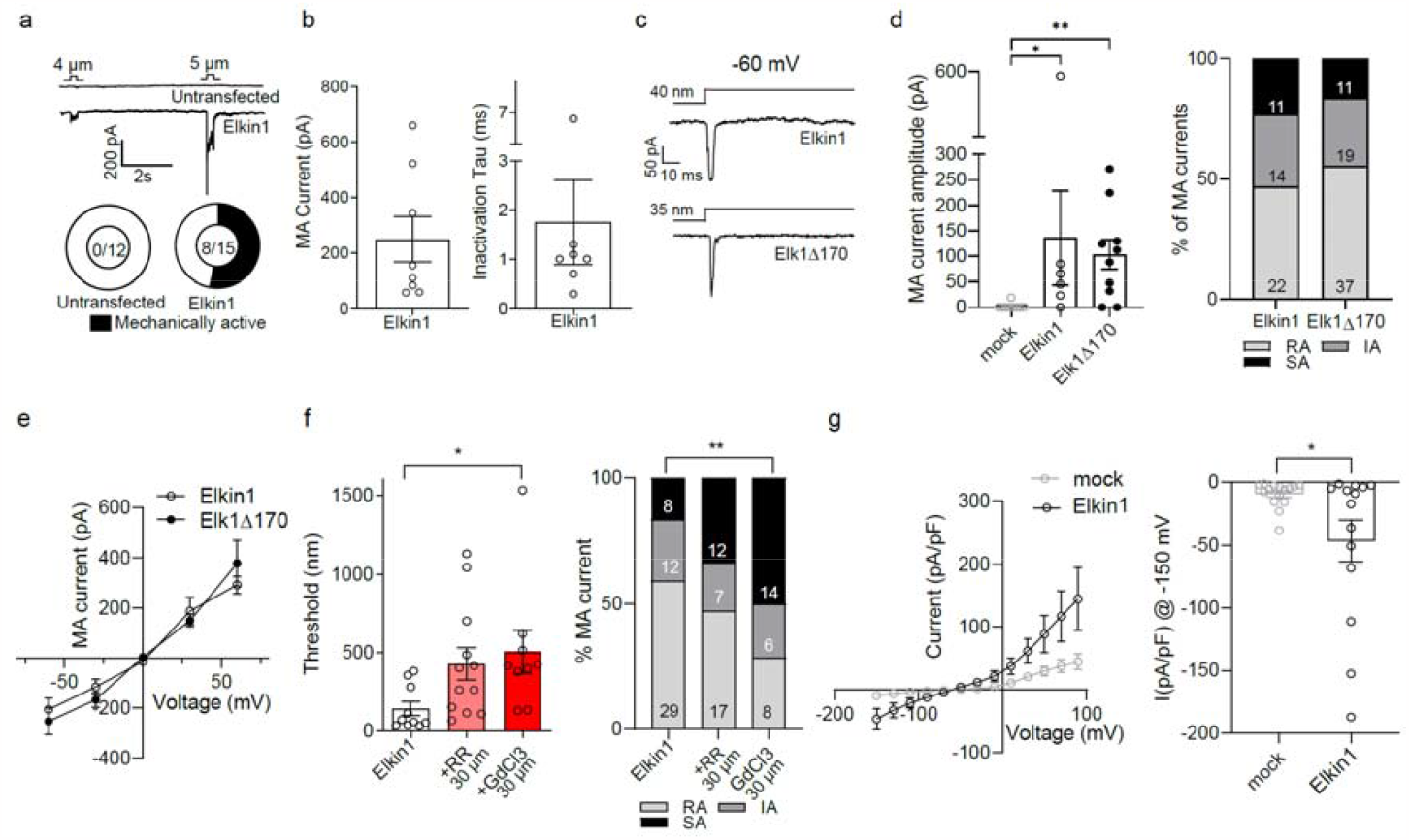
Elkin1 forms a mechanically-gated channel. **a**. Pie chart shows total number of mechanically active transfected cells. **a**,**b**. Poking-induced currents in HEK293T^*Piezo1-/-*^ cells upon transfection with mouse *Elkin1* cDNA. Dots represent individual cells. **c**. Representative MA-currents evoked by pillar deflection at -60 mV along with **d**. quantification of MA current amplitude and properties at 100-250 nm force bin (each dot represents a cell and numbers in bars are number of MA pillar stimuli) **e**. Current-voltage relationship of MA-currents evoked in cells transfected with *Elkin1* or *Elk1Δ170*. Dots are mean of n=10 in *Elkin1* and n=15 in *Elk1Δ170* transfected cells. **f**. Quantification of pillar deflection threshold and properties of Elkin1-dependent currents in the presence of pore blockers ruthenium red (RR) and GdCl_3_. Each dot represents a cell and numbers in bars are number of MA pillar stimuli. **g**. Current-voltage relationship of *Elkin1* transfected cells (each dot is a mean of n=14 mock and n=15 Elkin1 cells) as assessed through a series of voltage steps from -150 to 90 mV shows leak current properties of *Elkin1* transfected cells at -150 mV. Three group comparisons were made with one-way ANOVA followed by multiple comparison test and two group comparisons were made with Student’s t-test. Proportions were compared using chi-sq test. * indicates p < 0.05, ** indicates p < 0.01. Error bars = s.e.m.

Since Elkin1 is a mechanically-gated channel with properties similar to Piezo2 channels, we hypothesized that this protein could also be involved in mammalian touch sensation. We generated a CRISPR/Cas9 mediated genomic deletion of the mouse *Tmem87a/Elkin1* gene locus spanning sequences coding for transmembrane domains 2-6, that includes the proposed ion conduction pathway (Extended Data Fig. 2a)^14^. Mice homozygous for the genomic deletion (*Elkin1*^*-/-*^ mice) were viable and were born at the expected Mendelian ratios (WT: 25.7%, *Elkin1*^*+/-*^: 46.2%, *Elkin1*^*-/-*^: 28% n=132). Single molecule fluorescent *in situ* hybridisation (smFISH) and immunohistochemistry with an antibody against Elkin1 showed that *Elkin1*^*-/-*^ mice were complete null mutants (Fig 2a). Our validated Elkin1 antibody revealed that Elkin1 protein expression appeared to be especially high in sensory neurons of the DRG. Elkin1 protein was robustly detected in all subsets of DRG neurons, consistent with single cell expression data from mice, macaques and humans^18–22^, with ∼60% of neurons showing particularly high Elkin1 levels (Extended Data Fig. 2b,c). Sensory neurons expressing high levels of Elkin1 made up 34% of neurofilament heavy chain positive (NF200+) large neurons with myelinated axons. High levels of Elkin1 were also found in 75% of isolectin-B4 positive (IB4^+^) non-peptidergic small neurons^23^ and 45% of small neurons positive for the capsaicin-gated Transient receptor potential channel, Trpv1 (Extended Data Fig. 2b,c). Both of these two neurochemically distinct nociceptor types are reported to be responsive to mechanical force^24^.

**Figure 2:**
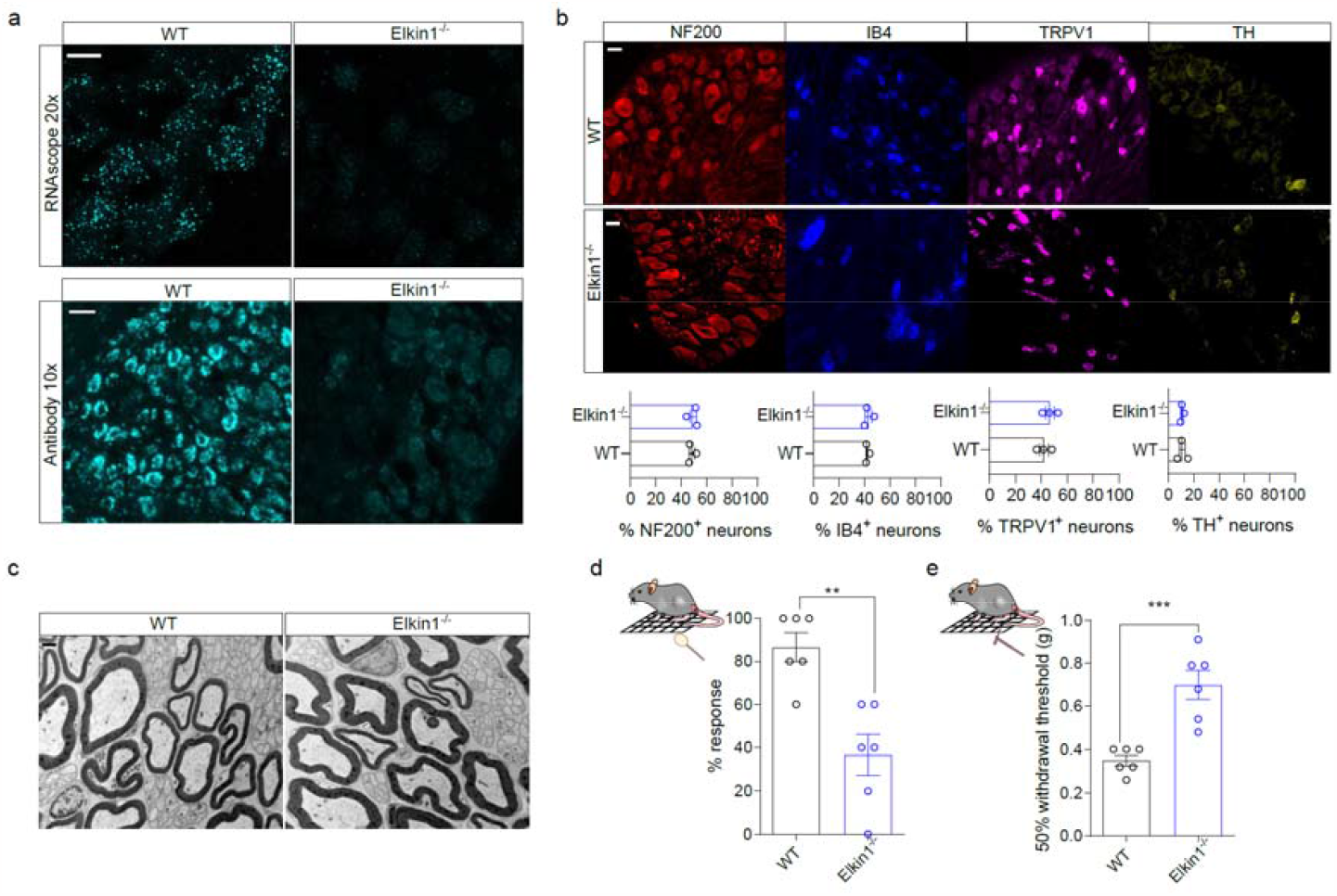
*Elkin1*^*-/-*^ mice are touch insensitive. **A**. Representative images of Elkin1 expression pattern, using a smFISH probe against *Elkin1* (top panel, scale = 20 μm) and antibody staining against Elkin1 (bottom panel, scale = 50 μm) from WT and *Elkin1*^*-/-*^ DRGs. **b**. Representative images of NF200 (red), IB4 (blue), TRPV1 (magenta) and TH (yellow) staining in WT and *Elkin1*^*-/-*^ DRGs (top panel) and quantification of percent of positive neurons in each group from 3 male mice. >500 neurons counted in each category. **c**. Ultra-structural analysis of saphenous nerve showed no deficit between WT and *Elkin1*^*-/-*^ mice. Scale bar = 1μm. **d:** Percent response of WT and *Elkin1*^*-/-*^ mice to brushing of a cotton swab **e**. Paw withdrawal threshold of WT and *Elkin1*^*-/-*^ mice to von Frey filaments. Two group comparisons were made with Student’s t-test. ** indicates p < 0.01, *** indicates p < 0.001. Error bars = s.e.m. Data from both male and female mice.

The absence of Elkin1 did not alter the neurochemical make-up of the sensory ganglia as the percentage of DRG neurons positive for markers like NF200, IB4, Trpv1 and tyrosine hydroxylase (TH) was unchanged in the *Elkin1*^*-/-*^ mice compared to wild types (WT) (Fig. 2b). Ultrastructural analysis of the saphenous nerve revealed no pathology or loss of myelinated or unmyelinated axons in *Elkin1*^*-/-*^ mice (Fig. 2c, Extended data table 1). However, behavioural indicators of touch sensitivity, like percentage of responses to a cotton swab, were profoundly reduced in *Elkin1*^*-/-*^ mice compared to WT (WT: 86.6% vs. *Elkin1*^*-/-*^: 36.6% paw withdrawal, Fig 2d). Paw withdrawal thresholds to von Frey filaments were also significantly elevated in *Elkin1*^*-/-*^ mice (Fig 2e), confirming reduced sensitivity to mechanical forces in *Elkin1*^*-/-*^ mice. However, non-mechanosensory modalities, like heat withdrawal thresholds, were unaltered in *Elkin1*^*-/-*^ mice (Extended Data Fig. 2d). *Elkin1*^*-/-*^ mice also showed no deficits in open field locomotion (Extended Data Fig 2e).

Large sensory neurons of the DRG are predominantly mechanoreceptors required for touch^8,12^. We therefore recorded MA-currents from isolated sensory neurons evoked by mechanical indentation and substrate deflection (Fig 3a, d)^8,25^. Normally, almost all large neurons exhibit robust predominantly RA MA-currents to both cell indentation and substrate deflection,^17,25^ which we confirmed here (Fig. 3a-f). However, only half of the large neurons (diameter >30 μm, Extended data Fig. 3a) from *Elkin1*^*-/-*^ mice displayed any MA-current (Fig. 3b,e). This insensitivity to mechanical stimuli was accompanied by a loss of RA MA-currents in *Elkin1*^*-/-*^ mice (Fig 3c, f). The current amplitude of MA-currents in the remaining mechanosensitive neurons was similar in WT and *Elkin1*^*-/-*^ mice, however, there was a small but significant increase in the deflection threshold in neurons from *Elkin1*^*-/-*^ mice (Extended data Fig. 3b,c). Large sensory neurons recorded from *Elkin1*^*-/-*^ mice also showed a slightly depolarized resting membrane potential consistent with the idea that this channel contributes to membrane leak (Extended data Fig. 3d). Strikingly, a significant loss of MA-currents was even detectable after loss of just one *Elkin1* allele (Extended data Fig. 3e). To show that the loss of MA-currents was not due to indirect effects of *Elkin1* gene inactivation, we conducted an acute rescue experiment. Using an adeno-associated virus neurotropic for sensory neurons (AAV-PHP.S-hSyn-dtom-mElkin1) we reintroduced Elkin1 back into acutely isolated sensory neurons from *Elkin1*^*-/-*^ mice. Elkin1 protein was detected in infected sensory neurons from *Elkin1*^*-/-*^ mice and there was a significant rescue of MA-currents - just 40% neurons had MA-currents in mock transfected cells compared to 75% of cells 48 hours after infection with AAV-PHP.S-hSyn-dtom-mElkin1 (Fig. 3g,h, Extended data Fig 4a).

**Figure 3:**
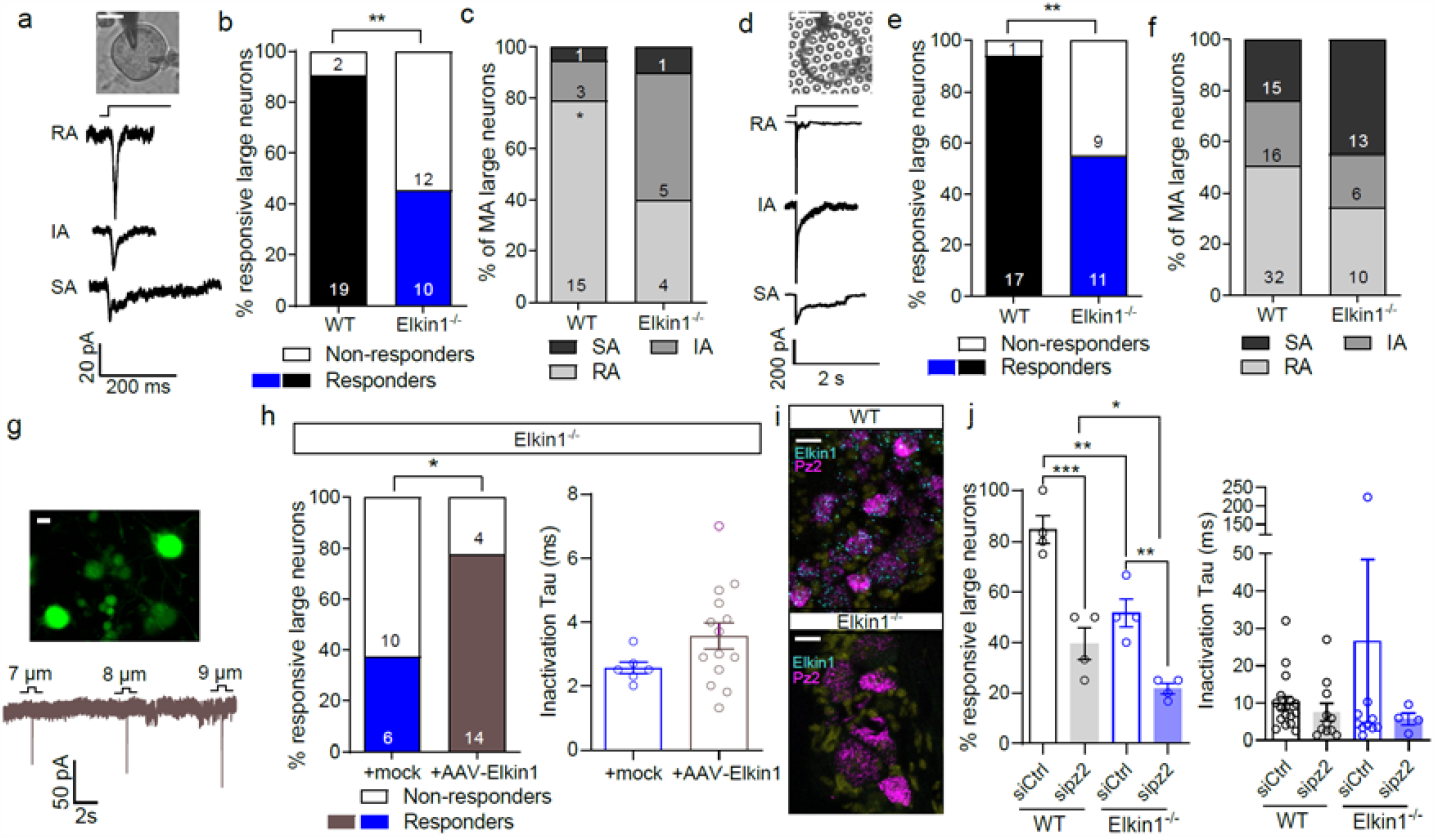
Elkin1 is necessary and sufficient for mechanically-gated currents in large DRG neurons. **a**. Representative traces of currents generated by large diameter neurons from of *Elkin1*^*-/-*^ mice. Scale bar = 20 μm. **b**. Percent of large diameter neurons with an MA-current in WT and *Elkin1*^*-/-*^ mice. **c**. Percent of rapidly, intermediate and slowly adapting, MA-currents in large diameter neurons. Number of cells are denoted in the bars. **d**. Representative traces of currents generated by pillar deflection of a large diameter neuron from a *Elkin1*^*-/-*^ mouse. Scale bar = 20 μm. **e**. Percent of mechanically sensitive large diameter neurons in WT and *Elkin1*^*-/-*^ mice in pillar assay. Number of cells are denoted in the bars. **f**. Percent of rapidly, intermediate and slowly adapting MA-currents in large diameter neurons. Number of MA pillar stimulations are denoted in the bars. **g**. top panel: Representative image of *Elkin1*^*-/-*^ DRG neurons stained by anti-RFP after being transduced by AAV-PHP.S-hSyn-dtom-mElkin1 (top panel, scale bar = 20 um). **g**. bottom panel: Representative traces of poking induced currents from a transduced neuron. **h:** left panel: Percent of mechanically active large diameter neurons upon transduction with AAV-PHP.S-hSyn-eGFP (mock) or AAV-PHP.S-hSyn-dtom-mElkin1.**h**. right panel: quantification of inactivation time constants of the measured currents. Each data point represents a single cell measured. **i**. Representative images of mRNA staining of whole DRG using probes against Piezo2 (*Pz2*, magenta) and Elkin1 (cyan). Scale bar = 50μm. **j**, left) Percentage of MA large neurons in WT or *Elkin*^*-/-*^ mice transfected with Control scrambled (siCtrl) or Piezo2 siRNA(sipz2). Each dot represents a mouse (number of neurons in each group is 20, 22, 19 and 19 respectively). **j**. right) Inactivation time constants of currents measured in each group. Proportions were compared using chi-sq test, four group comparisons were made using ANOVA followed by multiple comparison test. * indicates p < 0.05, ** indicates p < 0.01, *** indicates p < 0.001. Error bars = s.e.m. Data from both male and female mice.

The phenotype we observed in *Elkin1*^*-/-*^ sensory neurons was similar to the knockdown or genetic ablation of the Piezo2 mechanosensitive ion channel^3,5,26^. Using smFISH we detected colocalization of *Elkin1* and *Piezo2* mRNA in WT DRG neurons, but no change in *Piezo2* mRNA levels in *Elkin1*^*-/-*^ sensory neurons was observed (Fig. 3i, Extended data Fig. 4b). This suggests that ablation of *Elkin1* does not affect *Piezo2* expression. Next, we investigated whether the function of Elkin1 is dependent on Piezo2. Like others^3^, we could show that in WT DRG neurons, siRNA mediated knockdown of Piezo2 approximately halved the number of neurons with MA-currents (Fig 3j). If *Elkin1* exerts its function via *Piezo2*, knockdown of *Piezo2* in *Elkin1*^*-/-*^ neurons should not cause a further decrease in MA-currents. In contrast, we saw that MA-currents in *Elkin1*^*-/-*^ neurons could be reduced significantly further following *Piezo2* knockdown (Fig. 3j). Thus, neurons retaining MA-currents in *Elkin1*^*-/-*^ mice appear to have PIEZO2-dependent MA-currents.

To further test the idea that Elkin1 and Piezo2 function independently of each other we transfected of N2a^*Piezo1-/-*^ cells with expression constructs for *Piezo2* or double transfected with *Piezo2* and *Elkin1* and found that MA-currents were similar between single or double-transfected cells (Extended data Fig 4c). Thus, we found no evidence that Elkin1 was acting to modulate or sensitize Piezo2 channels. A known Piezo2 modulator is the MEC-2 related mechanotransduction protein Stoml3 which has been shown to dramatically sensitize PIEZO2 channels to substrate deflection^8,9,17,27^. We next asked whether there is also a molecular interaction between Stoml3 and Elkin1. Using a tripartite-GFP based protein complementation assay, we observed a robust green fluorescent signal indicating close association between Stoml3 and Elkin1 (Extended data Fig 5a). The protein complementation signal for a Stoml3/Elkin1 interaction was also blocked in the presence of the Stoml3 oligomerization blocker OB1 (Extended data Fig 5a)^9^. Interestingly, co-expression of *Stoml3* with *Elkin1* in HEK293T^*Piezo1-/-*^ cells revealed that Elkin1-dependent MA-currents displayed decreased mechanical thresholds and increased current amplitude in the presence of Stoml3 (Extended data Fig. 5b).

We next investigated whether Elkin1 is required for the mechanosensory function of identified mouse mechanoreceptors. First, we used an electrical search technique in an *ex vivo* saphenous skin nerve to functionally identify axons with and without mechanosensitvity^4,5,8,28^. Normally, single identified Aβ-fibres with the fastest conduction velocities (>10 m/s) always have a mechanosensitive receptive field as confirmed here for WT mice (Fig. 4a). However, blinded recordings made from *Elkin1*^*-/-*^ mice revealed that 40% of the Aβ-fibres had no detectable mechanosensitive receptive field (9/26 Aβ-fibres) (Fig 4a). We went on to examine the stimulus response properties of the remaining identified mechanoreceptors in the hairy skin. Aβ-fibre LTMRs innervating Merkel cells are classified as slowly-adapting mechanoreceptors (SAMs) with a dynamic and static response to ramp and hold force stimuli (Fig 4b). Normally around 50% of Aβ-fibres are classified as SAMs in the hairy skin^8,29^ and this was the case here in both WT and *Elkin1*^*-/-*^ mice (Extended Data Fig. 6a). However, the firing rates of SAMs to the static constant force phase of the stimulus was dramatically reduced at all stimulus strengths in *Elkin1*^*-/-*^ mice compared to controls (Fig. 4b). Indeed, a plot of the peristimulus time histogram for SAMs stimulated with 150 mN of force reveals that firing rates go down to almost zero just 3 seconds into a 10 second stimulus (Fig. 4c). However, the same SAMs from WT and *Elkin1*^*-/-*^ mice showed similar dynamic phase responses (Extended Data Fig. 6c). The remaining Aβ-mechanoreceptors were classified as rapidly adapting mechanoreceptors (RAMs) which only respond to skin movement and code the velocity of skin movement^11,12,30^. As a population, RAMs still coded the velocity of ramp stimuli but the overall firing rates were significantly lower in *Elkin1*^*-/-*^ mice compared to controls (Fig. 4d). These results could easily reflect loss of MA-currents in mechanoreceptors, but could also be due to morphological disruption of sensory endings. However, an analysis of mechanoreceptor endings in the skin of *Elkin1*^*-/-*^ mice did not reveal any obvious deficits (Extended data Fig. 6b). These results show that around half of the LTMRs are insensitive to mechanical forces in *Elkin1*^*-/-*^ mice, but also the remaining LTMRs showed profound functional deficits in their ability to detect mechanical force.

**Figure 4.**
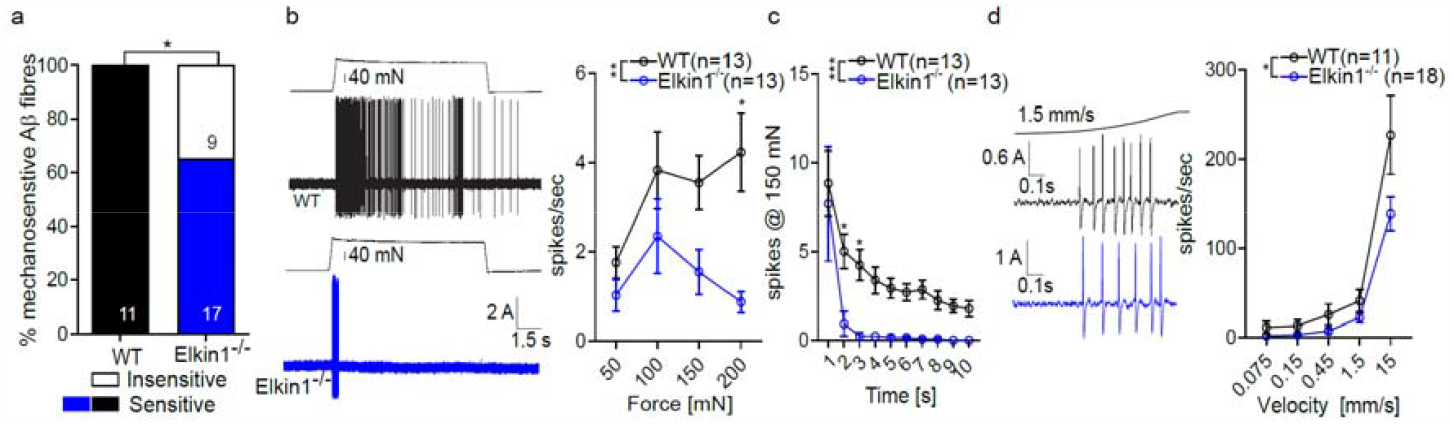
Elkin1 is required for LTMR function. **a**. Percent of mechanosensitive fast conducting, Aβ-fibres in the saphenous nerve assessed with an electrical search protocol **b**. Representative spikes evoked from slowly adapting mechanosensitive Aβ-fibres in WT and *Elkin1*^*-/-*^ mice and quantification of mean spike rates with increasing force. **c**. Absolute number of spikes over a 10s time period in the 150 mN force bin (each dot represents average response from fibres). **d**. Representative spikes evoked from rapidly-adapting mechanoreceptors to a moving ramp stimulus and quantification of the mean firing rates to ramps of increasing speed. Proportions were compared using chi-sq test (**a**.). All other group comparisons were made using 2 way ANOVA followed by multiple comparison. * indicates p < 0.05, ** indicates p < 0.01, *** indicates p < 0.001. Error bars = s.e.m. Data from at least 5 male and female mice.

Next, we made recordings from single mechanosensitive nociceptors in the saphenous nerve. Sensory neurons with thinly myelinated Aδ axons can be classified as A-fibre mechanonociceptors (AMs) that signal fast pain^31^ or D-hair receptors which are specialized LTMRs with directional sensitivity^32,33^. However, we found no change in the stimulus properties of D-hair or AM afferents in *Elkin1*^*-/-*^ mice compared to controls (Extended Data Fig. 6d). Many DRG with high levels of Elkin1 also appear to be nociceptors based on the presence of markers like IB4 and TRPV1^23^. We thus made a focused analysis of MA-currents in small/medium diameter neurons that displayed broad humped action potentials characteristic of nociceptors^23,25,34^ (Fig. 5a, Extended data Fig. 6). Many nociceptors lack MA-currents to cell indentation (∼40%)^24^, but this was not different in neurons recorded from *Elkin1*^*-/-*^ mice (Fig. 5 a,b). However, when the MA-currents were classified as RA (rapidly adapting, inactivation time <10 ms), IA (intermediate adapting, inactivation time constant 10-50 ms) and SA (slowly-adapting, inactivation time constant >50 ms) we saw a significant reduction in the proportion of RA MA-currents in *Elkin1*^*-/-*^ mice compared to WT (Fig. 5c). In addition, we saw a significant elevation in the amplitude of indentation needed to evoke the first MA-current in nociceptors from *Elkin1*^*-/-*^ mice (Fig. 5d). MA-currents evoked through substrate deflection showed no change between WT and *Elkin1*^*-/-*^ mice (Extended data Fig. 6b). We next focused our analysis on non-polymodal C-fibres that respond exclusively to mechanical stimuli and not to thermal stimuli as this population shows robust firing to mechanical force^35^. In the ramp and hold force protocol, mechanosensitive C-fibres from *Elkin1*^*-/-*^ mice showed slightly reduced firing across all stimulus strengths, but this did not reach statistical significance (Fig. 5e,f). We next analysed the time course of C-fibre activation during a 10 s long constant force stimulus. Normally C-fibres show a moderate degree of adaptation during constant force stimuli^5^. However, we found that even though initial firing rates were similar between genotypes at a stimulus strength of 100 mN firing rates dropped significantly more during the stimulus in C-fibres from *Elkin1*^*-/-*^ mice (Fig. 5g,f Extended data Fig. 8). Indeed reduced firing rates towards the end of the stimulus were observed for all intensities of stimulation (Extended data Fig. 8). Thus, the presence of Elkin1 in nociceptors appears to be necessary to maintain sensitivity constant forces that are known to be painful. This data suggest that Elkin1 may also be a relevant pharmacological target to treat mechanical pain.

**Figure 5:**
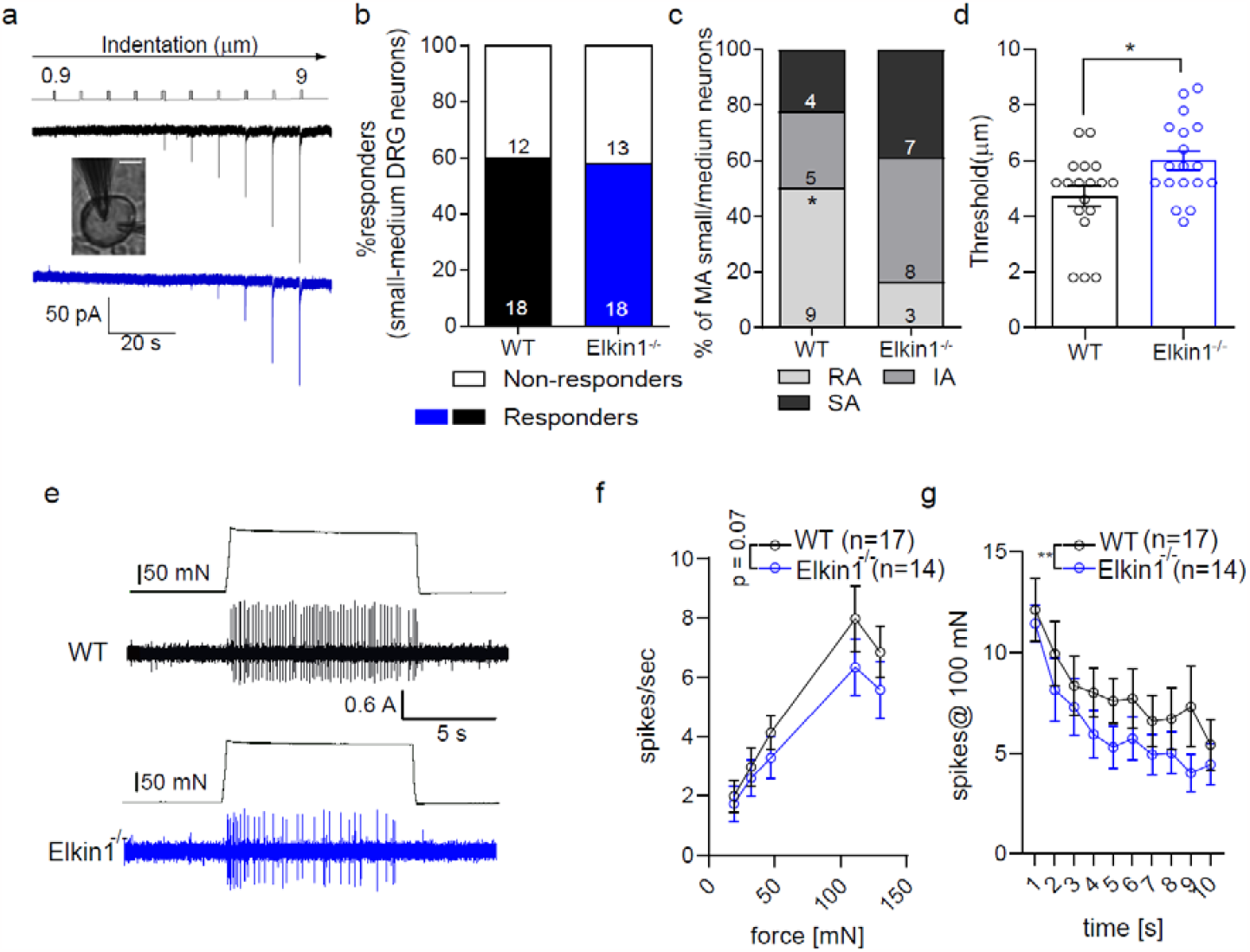
C-mechanonociceptors show reduced firing to sustained mechanical force in *Elkin1*^*-/-*^ mice. **a**. Representative poking current evoked from WT and a *Elkin1*^*-/-*^ small DRG neurons. Scale = 20 μm. **b**. Percent of mechanically sensitive small/medium diameter neurons in WT and *Elkin1*^*-/-*^ mice. **c**. Percent of rapidly, intermediate and slowly adapting MA-currents found in small/medium diameter neurons and their threshold for activation **d**. Number of cells are denoted as dots in the bar graph. **e**. Representative spikes from mechanosensitive C-fibres in WT and *Elkin1*^*-/-*^ mice. **f**. Quantification of the firing rates to increasing forces. **g**. Mean spiking rate over a 10s time period to a 100mN force stimulus Dots in **f**.,**g**. represent average of all fibres. Proportion was compared using chi-sq test. Two group comparison was made using Student’s t-test. All other group comparisons were made using 2-way ANOVA followed by multiple comparison. * indicates p < 0.05, ** indicates p < 0.01, *** indicates p < 0.001. Error bars = s.e.m. Data from at least 5 male and female mice.

Here we show that Elkin1 is absolutely necessary for the mechanosensory function of half of all LTMRs. *Elkin1*^*-/-*^ mice have electrically excitable sensory axons in the skin that were completely unable to respond to mechanical stimuli (Fig. 4a). The sustained firing of SAMs to constant force partly depends on Piezo2 expressed in mechanosensory Merkel cells^36^. Now our data suggest that Elkin1 is required for the Piezo2 independent transduction in SAMs as sustained responses were severely reduced in *Elkin1*^*-/-*^ mice (Fig. 4b,c). Consistent with the expression pattern of Elkin1, we noted that maintained firing of C-fibre nociceptors to constant force was also impaired in the absence of the Elkin1 protein. The loss of mechanically-gated currents, impaired touch driven behaviour and deficits in LTMR function are all features of mice lacking Stoml3 and features of conditional *Piezo2* mutants ^5,8,26,37^. Our data supports a model where Elkin1 and Piezo2 channels are used to preferentially support sensory mechanotransduction in distinct subpopulations of LTMRs, and that both can be modulated by Stoml3. There is evidence that Stoml3 can also modulate MA-currents in nociceptors consistent with a role for Elkin1 in conferring robustness to the C-fibre responses to force^17^. Here we identify a new mechanically gated-ion channel that is necessary for somatosensory function which will enable the identification of new molecular partners that can account for the entirety of touch transduction.

## Methods

### Animals

C57BL/6N adult (10-30 weeks) mice of both sexes were used in the study. Mice were housed in groups of 5 with food, water and enrichment available ad libitum. All animal protocols were regulated by the German federal authorities (State of Berlin). *Elkin1*^*-/-*^ mouse line was generated by the Ingenious Targeting Laboratory (Ronkonkoma, NY) using CrispR/Cas9 strategy by deletion of exons 8 through 15 introducing a frameshift as illustrated in Extended Fig 2a. The genotyping strategy utilized was as follows: For mutant genotyping: Forward primer: 5’-AAGGTGAAATAGCTCTCGATGG -3’, reverse primer: 5’-GGGAAAGAACGGGAACACA -3’ (expected band = 640 bp). For WT genotyping: Forward primer: 5’ -TTTGCCAGTGACTGTTGAAGT – 3’, Reverse primer: 5’ - AGATAAAGGGGTCCCACAGGA – 3’ (expected band = 481 bp).

### Behaviour

For all behavioural assays animals were habituated in the test paradigm for two days and analysis of the behaviours were carried out in a blinded manner.

Animals were placed on a raised platform with mesh-like floor (5x5 mm, von Frey apparatus) until they produced minimal movement. Paw withdrawal threshold was then assessed using a standard Semmes-Weinstein set of von Frey filaments (ranging from 0.07 - 2 g, Aesthesio) in accordance to the up-and-down method^38,39^. 50 % paw withdrawal threshold was calculated using the open source software, Up-Down Reader (https://bioapps.shinyapps.io/von_frey_app/)^39^.

Withdrawal response to cotton swab was assessed as a measure of evoked responses to very small stimuli (0.7 – 1.6 mN^40^), while evoked responses to brush was used to measure intermediate touch sensation (>4 mN). For these tests, after habituation on the von Frey apparatus, a cotton swab (puffed out to ∼ 3 times its original volume) or Number 0 brush was applied in a sweeping motion across alternate hind paws, for a total of 5 times. The number of withdrawals throughout the trials was counted and coded as percentage of total trials.

Hargreaves assay was used to assess the temperature sensitivity of mice. Briefly, mice were habituated on a plexiglass apparatus until minimal movement was observed. Their hind paws were then stimulated from below with an infrared light source (Ugo Basile, 37370) and the time required to withdraw the paws were recorded as indicators of their ability to assess temperature.

Male and female mice aged between 31-35 days were used to assess locomotor behaviour using the open field test. Testing was conducted during the inactive phase between 10 am and 3 pm. Before testing, the animals were habituated to the test room for at least 30 minutes.

The OFT apparatus consisted of an open arena (50x50 cm) where mice were allowed to freely explore for 10 minutes. Animal locomotion and location were recorded and tracked automatically using the ActiMot2 system (TSE Systems GmbH, Germany). Additionally, all trials were recorded using video cameras for subsequent detailed analyses. Between each trial, the arenas were cleaned using 80% ethanol and water and briefly dried with paper towels.

### DRG neuron culture

Lumbar DRGs (L2-L5) were collected from mice into plating medium (DMEM-F12 (Gibco) supplemented with 10% fetal horse serum (FHS, Life Technologies) and 1% penicillin and streptomycin (P/S, Sigma-Aldrich)). These DRGs then underwent enzymatic digestion in 1.25% Collagenase IV (1 mg/ml, Sigma-Aldrich) for 1 hour at 37°C and 2.5% Trypsin (Sigma-Aldrich) for 15 min, at 37°C, followed by mechanical trituration with a P1000 pipette tip and purification in a 15% fraction V BSA column. The neurons were plated on glass bottomed plates (for indentation assays) or elastomeric pillar arrays (made as descried before^17^) coated with poly-L-lysine and laminin. Neurons were cultured overnight in an incubator (37°C and 5% CO2) and electrophysiological experiments were performed 18-24 hours after plating.

### AAV transduction

AAV-PHP.S-hSyn-dtom-mElkin1 (4.88 × 10^12^ vg/ml, canonical form of Elkin1 Uniprot ID: Q8NBN3-1) and AAV-PHP.S-hSyn-eGFP (4.94 × 10^12^ vg/ml) were manufactured in the Charité Viral Core facility (Berlin, Germany). For viral transduction of DRG neurons, titre-matched number of viruses were mixed with dissociated DRG neurons (after the mechanical trituration step described above) and plated on glass-bottomed dishes (50 μl per dish) coated with poly-L-lysine and laminin. Neurons were allowed to attach to plates for 2 hours before being submerged by plating medium. Some of the plated neurons were fixed and immunocytochemistry was performed using anti-Tmem87a (host: rabbit, HPA018104, 1:100) and anti-RFP (host: rabbit, AB34771, 1:500) antibodies to quantify percent of viral transduction.

### siRNA transfection

2 hours after plating the DRG neurons in glass bottomed dishes, siRNA transfection was carried out using DharmaFECT (Horizon) reagents according to the manufacturers’ guidelines. Briefly, siRNAs were mixed with serum and antibiotics free DRG neuron plating medium in a tube (total volume of 100 μl per dish), and 4.5 μl of DharmaFECT 4 transfection reagent was mixed with serum and antibiotics free DRG neuron plating medium (total volume of 100 ul) in another tube. Each tube was incubated separately for 5 min at RT, then mixed together and incubated for a further 20 min. 800 μl of antibiotics free complete medium was added to this mixture and added to the neurons. Experimental dishes were transfected with a final concentration of 50 nM ON-TARGETplus SMARTpool mouse Piezo 2 and 50 nM of siGLO green transfection indicator and control dishes were transfected with 50 nM of ON-TARGETplus Non-targeting Pool and 50 nM of siGLO green transfection indicator. Neurons were patched 48 hours after transfection with siRNA.

### Whole cell electrophysiology

Whole-cell patch clamp experiments were performed using heat-polished borosilicate glass pipettes (Harvard apparatus, 1.17 mm x 0.87 mm) with a resistance of 3-6 MΩ. The pipettes were pulled using a DMZ puller (Germany) and filled with a solution containing (in mM): 110 KCl, 10 NaCl, 1 MgCl_2_, 1 EGTA and 10 HEPES, pH adjusted to 7.3 with KOH. Current-clamp experiments were performed to classify sensory neurons into mechano- and nociceptors. The extracellular solution contained (in mM): 140 NaCl, 4 KCl, 2 CaCl_2_, 1 MgCl_2_, 4 Glucose and 10 HEPES, pH adjusted to 7.4 with NaOH.

Currents were evoked either using pillar deflection or indentation of the soma (poking assay) at a holding potential of -60 mV. For pillar arrays experiments, a single pillar was deflected using a heat-polished borosilicate glass pipette (mechanical stimulator) driven by a MM3A micromanipulator (Kleindiek Nanotechnik, Germany) as described previously. Bright field images (Zeiss 200 inverted microscope) were collected using a 40X objective and a CoolSnapEZ camera (Photometrics, Tucson, AZ) before and after the pillar stimuli to calculate the pillar deflection. The light intensity of the center of each pillar before and after the stimuli was used to fit a 2D-Gaussian (Igor Software, WaveMetrics, USA) in order to calculate the pillar displacement. For mechanosensitive current-voltage curve with HEK293T^Piezo1-/-^ cells, pillar displacements were produced at -60, -30, 0, 30 and 60 mV. For poking experiments, indentation was produced on the neuronal cell membrane within the range of 1 – 9 μm using a heat-polished borosilicate glass pipette (mechanical stimulator) driven by a MM3A micromanipulator. For experiments with antagonists, cells were bathed in 30 μm ruthenium red (Sigma, 557450, diluted in ECS from a stock of 10 mM in water) and 30 μm GdCl_3_ (Tocris, 4741, diluted in ECS from a stock of 100 mM in water).

Currents and the biophysical parameters were analysed using FitMaster (HEKA, Elektronik GmbH, Germany).

### Ex vivo skin nerve

Ex vivo skin nerve electrophysiology from cutaneous sensory fibres of the saphenous nerve was conducted following the method previously described^32^. Briefly, mice were euthanized by CO_2_ inhalation for 2-5 min followed by cervical dislocation, and the hair of the limb was shaved off. The hairy skin from the hind paw was removed along with the saphenous nerve up to the hip.

The innervated hairy skin was transferred to a bath chamber which was constantly perfused with warm (32°C) carbonated (95% O2, 5% CO2) interstitial fluid (SIF buffer): 123 mM NaCl, 3.5 mM KCl, 0.7mM MgSO4, 1.7 mM NaH2PO4, 2.0 mM CaCl2, 9.5 mM sodium gluconate, 5.5 mM glucose, 7.5 mM sucrose and 10 mM HEPES (pH 7.4). The skin was pinned out and stretched, such that the inside of the skin could be stimulated using stimulator probes. The nerve was fed through a small opening to an adjacent chamber filled with the mineral oil, where fine filaments were teased from the nerve and placed on a silver wire recording electrode.

Mechanically sensitive units were first located using blunt stimuli applied with a glass rod. The spike pattern and the sensitivity to stimulus velocity were used to classify mechanoreceptors as previously described^8,9,32^. The raw electrophysiological data were recorded using a Powerlab 4/30 system and Labchart 8 software with the spike-histogram extension supported by an oscilloscope for visual identification and later analysis. All mechanical responses analyzed were corrected for the latency delay between the electrical stimulus and the arrival of the action potential at the electrode. The conduction velocity (CV) was measured with the formula CV = distance/ time delay, in which CVs > 10 ms^−1^ were classified as RAMs or SAMs (Aβ, <10 ms^−1^ as Aδ and <1.5 ms^−1^ as C-fibers).

Mechanically sensitive units were stimulated with the use of a piezo actuator (Physik Instrumente (PI) GmbH & Co. KG, model P-841.60) connected to a force sensor with calibrated conversion factor of Volts to Milinewtons. Different mechanical stimulation protocols were used to identify and characterize the sensory afferents. All types of mechanoreceptors were tested with a vibrating stimulus with increasing amplitude and 20 Hz frequency. The force needed to evoke the first action potential was measured. Additionally, a ramp and hold step was used with constant force (100 mN) and repeated with varying probe movement velocity (0.075, 0.15, 0.45, 1.5, and 15 mm s−1). Only the firing activity evoked during the dynamic phase was analyzed. SAM mechanoreceptors, Aδ mechanoreceptors, and nociceptors were tested mechanically with a constant ramp (1.5–2 mN ms^−1^) and hold (10 seconds of static phase) stimulation. Force of increasing amplitude was applied in five consecutive indentations, spikes evoked during the static phase were analyzed.

### Cell line culture

HEK293T^*Piezo1-/-*^ and N2a^*Piezo1-/-*^ cells were used for characterisation of the currents evoked by mElkin1. HEK293T^Piezo1-/-^ were cultured in DMEM-Glutamax (gibco, ThermoFisher scientific) supplemented with 10% fetal bovine serum (FBS, PAN Biotech Gmbh) and 1% penicillin and streptomycin (P/S, Sigma-Aldrich). N2a^*Piezo1-/-*^ cells were cultured in DMEM-Glutamax (gibco, ThermoFisher scientific) supplemented with 45% Opti-MEM (gibco, ThermoFisher scientific), 10% FBS and 1% P/S. HEK293T^*Piezo1-/-*^ and N2a^*Piezo1-/-*^ cells were transiently transfected with 4 μl FuGene6 per 100 μl of Opti-MEM with 1 μg of cDNA as per the manufacturer’s instructions. Cells were maintained in serum free medium overnight before electrophysiological recordings in a 37°C and 5% CO2 incubator. Data were obtained from at least 3 transfections. Plasmids used for transfection in electrophysiological studies are as follows: mPiezo2-IRES-EGFP, mElkin1-IRES-dtomato, mElkin1-IRES-EGFP, mElkin1-Δ1-170-IRES-EGFP, mStoml3-2A-tdTomato. All mElkin1 are Uniprot ID: Q8NBN3-1.

### Immunohistochemistry

L2–L5 DRG were collected from animals and postfixed in Zamboni’s fixative for 1 hour, followed by overnight incubation in 30% (w/v) sucrose (in PBS) at 4°C for cryoprotection. DRG were next embedded in Shandon M-1 Embedding Matrix (Thermo Fisher Scientific), snap frozen on dry ice and stored at −80□°C. Embedded DRG were sectioned (12□μm) using a Leica Cryostat (CM3000; Nussloch), mounted on Superfrost Plus microscope slides (Thermo Fisher Scientific) and stored at −80□°C until staining. During staining, slides were washed with PBS-tween and blocked in antibody diluent solution: 0.2% (v/v) Triton X-100, 5% (v/v) donkey serum and 1% (v/v) bovine serum albumin in PBS for 1 hour at room temperature before overnight incubation at 4□°C with primary antibodies. The following primary antibodies were used on DRG tissue slides: anti-Tmem87a (host: rabbit, HPA018104, 1:100), anti-tyrosine hydroxylase (TH, host: sheep, AB1542, 1:1000), anti-NF200 (host: chicken, AB72996, 1:1000), anti-Trpv1 (host: goat, SC14298, 1:200).

Slides were then washed three times using PBS-tween and incubated with the following species-specific conjugated secondary antibodies at 1:500 - Alexa Fluor 488 anti-rabbit (A21206), Alexa Fluor 568 anti-sheep (A21099), Alexa Fluor 568 anti-chicken (A11041), Alexa Fluor 555 anti-goat (A21432) or Isolectin GS B4 conjugated to Alexa 594 (121413, Thermo Fisher) for 2 hours at room temperature (20-22□°C). The secondary antibody was washed three times in PBStween, mounted with Dako (Agilent, S3023) and imaged with a Leica fluorescent microscope. Exposure levels were kept constant for each slide and the same contrast enhancements were made to all slides. Negative controls without the primary antibody showed no staining with either secondary. *Elkin1*^*-/-*^ mice did not show staining with anti-Tmem87a antibody. Positive neurons were scored as using an R toolkit (https://github.com/amapruns/Immunohistochemistry_Analysis) followed by manual validation. Briefly, mean gray value (intensity) of all neurons in 1–3 sections from each DRG level of interest in each mouse was measured using ImageJ. A neuron was scored as positive for a stain if it had an intensity value >2 SDs above the average normalized minimum gray value across all sections.

Free-floating immunohistochemistry was conducted on 40 μm sections of hairy skin as described above with minor adjustments such as longer washes and incubation in primary antibodies (anti-NF200 (host: chicken, AB72996, 1:500), anti-S100B (host: rabbit, 15146-1-AP, 1:200) for 72 hours and in secondary antibodies (Alexa Fluor 488 anti-chicken, A11039, 1:500, Alexa Fluor 568 anti-rabbit, A10042, 1:500) overnight. The free-floating sections were then mounted on glass slides using 0.2% gelatine in PBS and imaged using a ZEISS confocal microscope.

### SmFISH

*Elkin1* and *Piezo2* mRNA levels were assessed in DRGs from WT (n = 3) and *Elkin1*^*-/-*^ (n = 3) mice using RNAscope assay (a fluorescent in situ hybridization technology). Briefly, snap-frozen and embedded DRGs were sectioned and mounted on coverslips as described above. Next, hybridization of probes and subsequent signal development was carried out using H_2_O_2_ and Protease Reagents and Multiplex Fluorescent Detection Reagents v2 kits (Advanced Cell Diagnostics (ACD); Hayward, CA) as described in the manufacturer’s protocol. The following probes were used: Piezo2 - Mm-Piezo2-E43-E45 (C1 #439971) and Elkin1 - Mm-Tmem87a (C3 #868581). The fluorophores applied to detect the signal from the probes were Opal 690 Reagent (1:750; Akoya Biosciences, #FP1497001KT) for C1 and Opal 570 Reagent (1:750; Akoya Biosciences, #FP1488001KT) for C3. Lastly, slides were incubated with RNAscope DAPI and cover slipped using Fluorescence Mounting Medium (Dako)

### Electron microscopy

Saphenous nerves from WT and *Elkin1*^*-/-*^ mice were examined using a Zeiss 910 electron microscope. Briefly, nerves were isolated and fixed in 4% paraformaldehyde/2.5% glutaraldehyde in 0.1 M phosphate buffer for at least 24 hours at 4°C, before being treated with 1% OsO4 for 2 hours and dehydrated in an ethanol gradient and propylene oxide. Then, the nerves were embedded in polyresin, cut using a microtome in ultrathin sections (70 nm) ad imaged at 2500x magnification. Analysis of the A and C fibre dimensions were made^41^ using https://axondeepseg.readthedocs.io/en/latest/ followed by manual quality control by a blinded experimenter.

### Tripartite split-GFP complementation assay

The tripartite split-GFP complementation assay (modified from^42^) was performed in HEK293T (ACC 635, DSMZ). HEK293T cells were cultured in DMEM (DMEM 41966) supplemented with 10%FCS and 1% Penicillin/Streptomycin. HEK293T cells were cultured to about 70% confluence; cells were co-transfected, as indicated, with GFP1-9::iRFP702 (Addgene #130125), mouseStoml3-GFP10::mCHERRY and mouseElkin1-GFP11::eBP2 plasmids using Fugene HD and following manufacturer’s instructions (Promega). 7 hours after transfection, the culture medium was changed with fresh medium containing DMSO or OB1 (5μM). After 40 hours, cells were fixed in 4% PFA and images (eGFP, mCherry and eBFP2) were acquired at cytationC10 (Biotek, Objective 60x). eGFP (interaction signal) and mCherry (transfection control signal) % of cells and fluorescence intensity were analysed using Gen5 (Biotek). eGFP signal was normalized with mCherry signal (% eGFP/mCherry and fluorescence intensity eGFP/mCherry), data were then normalized to control (DMSO).

### Statistical analysis

All statistical tests used are detailed in the figure legends. Comparisons between two groups were carried out using Student’s t-test and more than three groups using ANOVA with post hoc tests. Proportions were compared using the chi-sq test. Electrophysiological skin nerve data consisting of two groups and multiple time points/forces were compared using 2-way ANOVA.

## Acknowledgement

This research was funded by an ERC grant to GRL (Sensational Tethers 789128). SC and AR were recipients of Alexander von Humboldt research fellowship. We thank Franziska Bartlett and Letizia Dalmasso for help with mouse genotyping, Bettina Purfürst for electron microscopy. We thank James Poulet, Stefan Lechner and members of the Lewin lab for constructive comments on the manuscript.

## Author contributions

Conceptualization S.C and G.R.L.; mouse model design: G.R.L and V.B.; Patch clamp physiology and anatomy and cell biology S.C with help from O.C-S, K.P. and Z.B. Antibody validation C.F, A.H.; Tripartite GFP design and implementation R.F, A.R and A.T-L. H, Molecular biology A.T-L.H. Skin nerve electrophysiology J.D.K (nociceptors), M.A.K (mechanoreceptors, A-mechanonociceptors D-hair) and G.R,L. (electrical search); Behavioural assessment S.C with help from T.P., H.H.; Electron microscopy and analysis J.D.K and S.C.; Writing S.C. and G.R.L with input from all authors; Supervision and funding: G.R.L.

The authors declare no conflict of interest

Materials and Correspondence should be addressed to G.R.L.

