## Supplementary Figures for "Touch sensation requires the mechanically-gated ion channel Elkin1"

|  | **WT (n = 3)** | **Elkin1^-/-^ (n = 3)** |
| --- | --- | --- |
| **Avg cross-sectional area of A-fibres** | 4.5 ± 0.2 | 4.4 ± 0.1 |
| **G ratio of A-fibres** | 0.69 ± 0.007 | 0.68 ± 0.008 |
| **Avg cross-sectional area of C-fibres** | 0.29 ± 0.05 | 0.29 ± 0.02 |
| **Total A-fibres (extrapolated)** | 565.0 ± 53.3 | 663.7 ± 104.9 |
| **Total C-fibres (extrapolated)** | 2935.0 ± 340.9 | 3108.0 ± 373.0 |
| **Ratio of A fibres to C fibres** | 5.3 ± 0.7 | 5.7 ± 1.1 |

**Extended Data Table 1: Anatomical properties of the saphenous nerve assessed using electron microscopy.**

Data table showing no change in ultra-structure of saphenous nerve in WT and *Elkin1^-/-^* mice. Data obtained from male mice, 15 images from each mice were analyzed. Data presented as mean ± SEM.

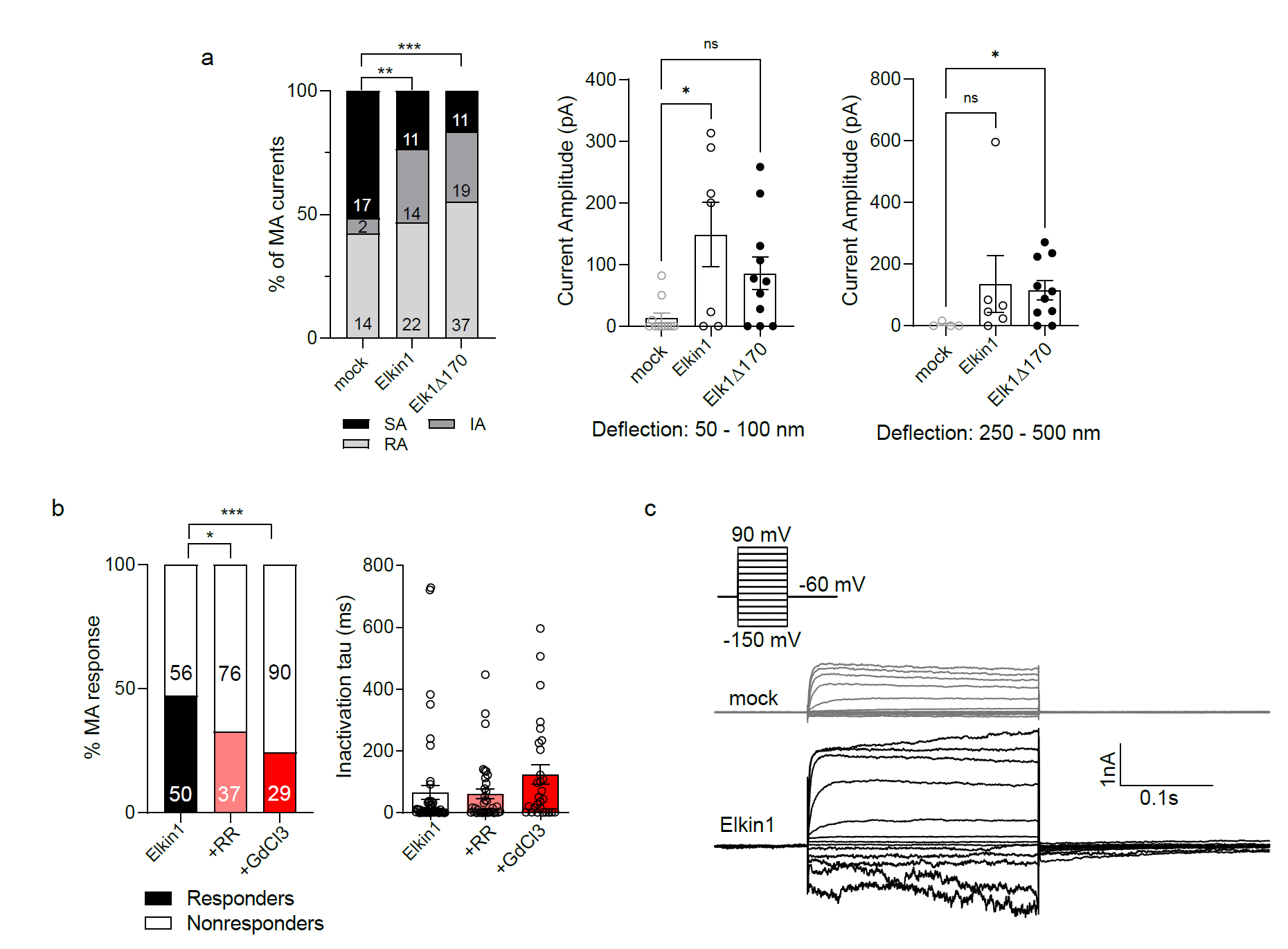

**Extended Data Fig 1: Biophysical properties of Elkin1.** a) (left) Percent of rapidly adapting (RA), intermediately adapting (IA) or slowly adapting (SA) MA currents evoked by pillar assay (substrate deflection) in HEK293T*^Piezo1-/-^* cells. Numbers in bars represent number of deflections. (right) Amplitudes of MA currents evoked at 50-100 nm and 250-500 nm force bins. b) Percent of mechanically active pillar stimulations and their inactivation time constants in Elkin1 transfected HEK293T*^Piezo1-/-^* cells when incubated with 30 µm ruthenium red (RR) and 30 µm GdCl_3_. Numbers in bars represent number of pillar stimulations. c) Raw traces of leak currents in mock (grey) and Elkin1 (black) transfected HEK293T*^Piezo1-/-^* cells. Proportions were compared using chi-sq tests, three group comparisons were made using ANOVA followed by multiple comparison tests. * indicates p < 0.05, ** indicates p < 0.01, *** indicates p < 0.001. Error bars = SEM.

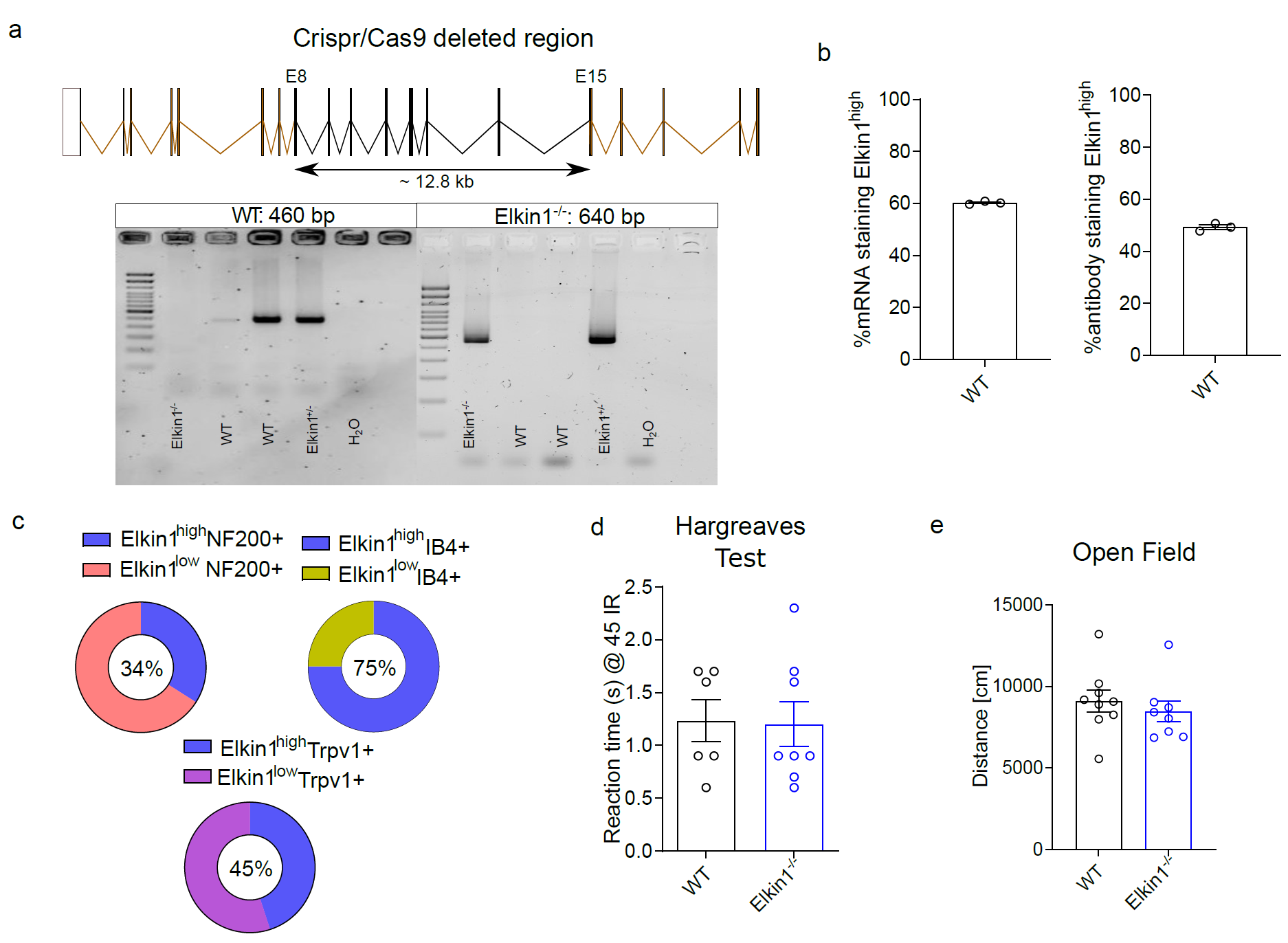

**Extended Data Fig 2: Characterization of Elkin1^-/-^ mice.** a) Representation of the strategy to knockout Elkin1 using Crispr/Cas9 along with PCR bands from genomic DNA of WT, Elkin1^+/-^ and Elkin1^-/-^ mice. b) Percent of DRG sensory neurons expressing Elkin1 at a high level as assessed through RNAscope (left) and immunohistochemistry (right). Dots represent each mouse, > 100 neurons were analyzed in each mouse. c) Quantification of colocalization between sensory neurons expressing high levels of Elkin1 and positive for NF200 (orange), IB4 (yellow) and Trpv1 (magenta). Number inside circles represent the highly expressing Elkin1 group. d) Hargreaves assay for thermal perception in WT and Elkin1^-/-^ mice. e) Open field test showing total distance travelled. Error bars = SEM.

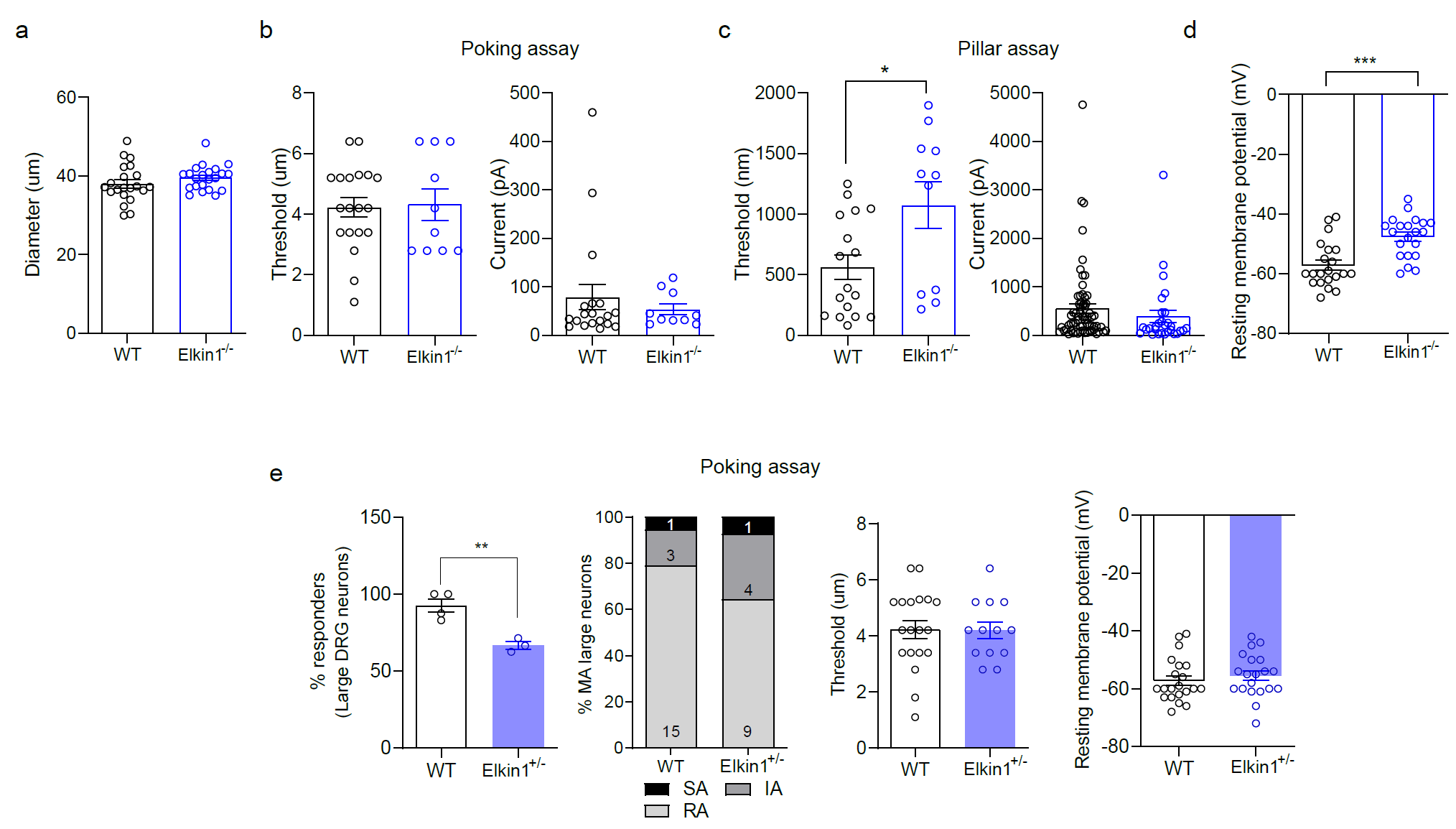

**Extended Data Fig 3: Electrophysiological properties of large neurons.** a) Diameter of neurons recorded and classified as large diameter neurons. b) Threshold of MA current activation in the poking assay along with the max amplitude of these currents. c) Threshold of MA current activation in the pillar assay (each dot represents individual cells) along with the max amplitude of these currents (each dot represents individual stimulation). d) resting membrane potential of the large neurons. e) Properties of MA currents in Elkin1+/- mice as assessed through poking assay including percent MA, percent of rapidly (RA), intermediately (IA) or slowly adapting (SA) MA currents, threshold of MA current activation and resting membrane potential. WT data is the same used for comparison with Elkin1^-/-^ mice. Data obtained from both male and female mice. Two groups were compared using two-sided Student’s t-test. ** indicates p < 0.01, *** indicates p < 0.001. Error bars = SEM.

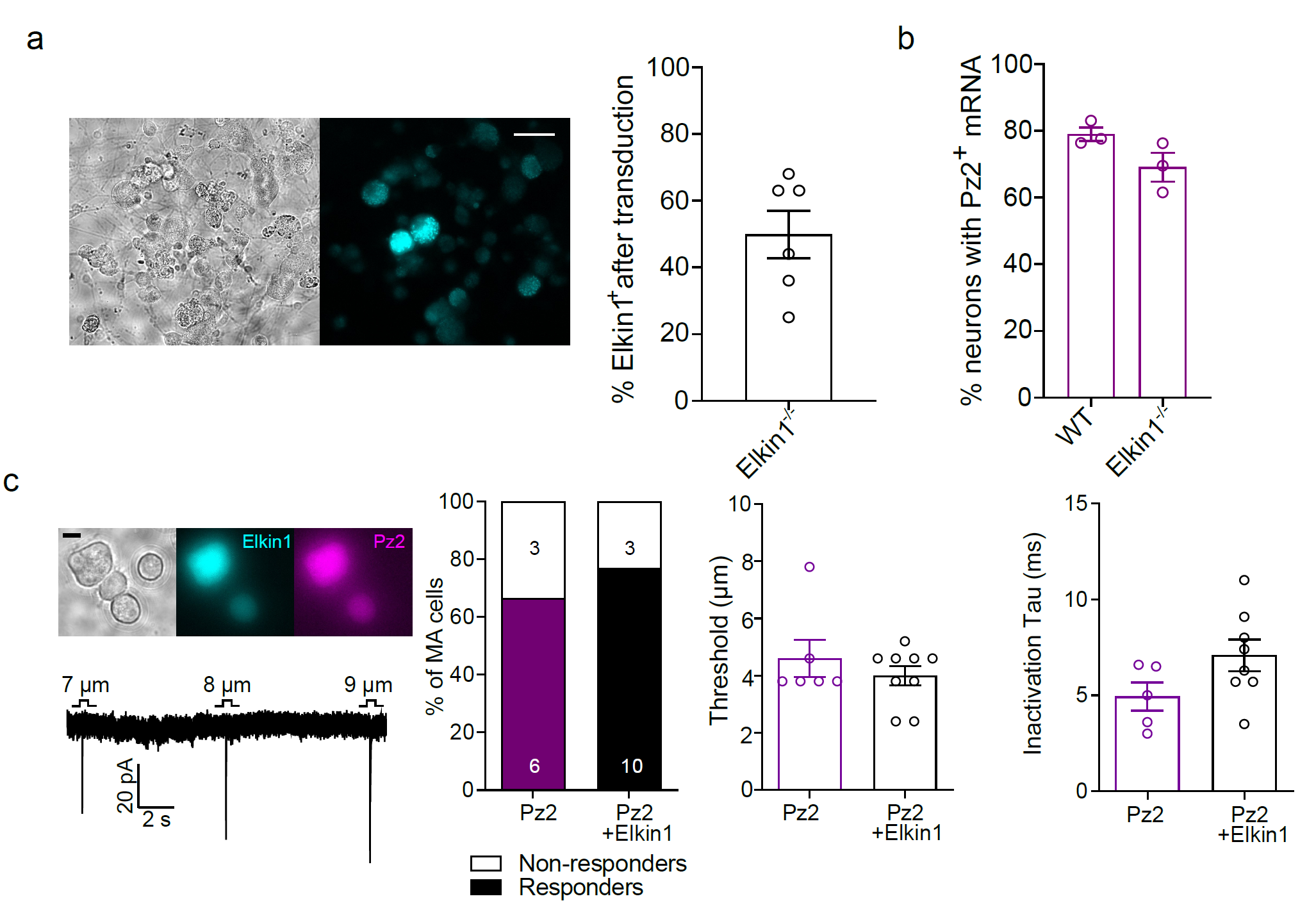

**Extended Data Fig 4: Elkin1 replacement and colocalization with Piezo2.** a, left) Representative image of sensory neurons from Elkin1^-/-^ mice transduced with AAV-PHP.S-hSyn-dtom-mElkin1-iso1 and stained with anti-Tmem87a (Elkin1) antibody. Scale = 100 µm. (right) quantification of the percent of Elkin1 positive neurons after viral transduction. Dots represent 2 dishes each from 3 mice. b) percentage of DRG neurons positive for Piezo2 mRNA as assessed through RNAscope – representative images in main Fig 2. c) N2a^Piezo1-/-^ cells transfected with Elkin1 and Piezo2 cDNA showing similar percent of MA cells, threshold of MA current activation and intactivation time constant as assessed through poking assay. Scale = 10 µm. Error bars = SEM.

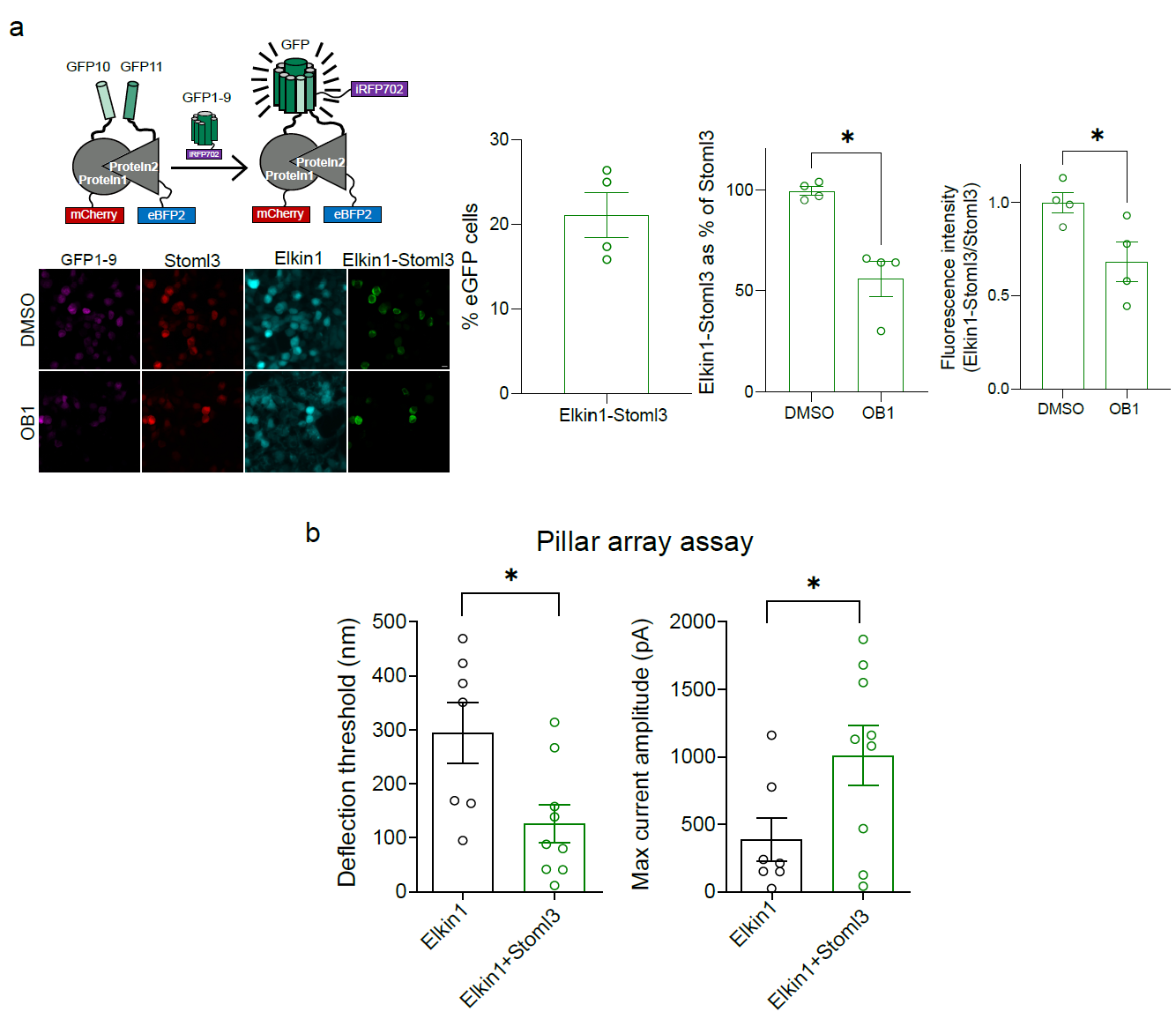

**Extended Data Fig 5: Stoml3-Elkin1 interactions.** a) Tripartite GFP based protein complementation assay showing percent interaction of Stoml3 and Elkin1 (% eGFP cells) along with decrease in this interaction when incubated with an antagonist of Stoml3 oligomerization, OB1. b) Threshold of MA current activation and peak MA current amplitude evoked by pillar deflections in HEK293T*^Piezo1-/-^* cells transfected with Elkin1 and Elkin1+Stoml3. Scale bar = 10 μm. Two groups were compared using two-sided Student’s t-test. * indicates p < 0.05. Error bars = SEM.

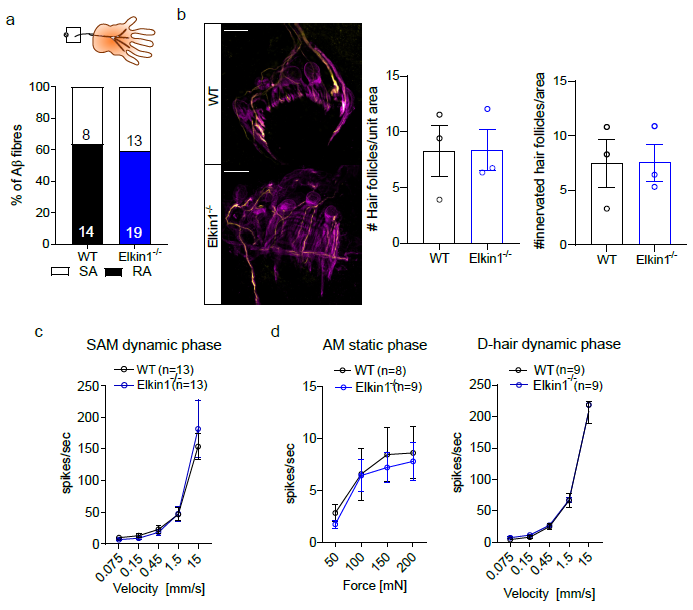

**Extended Data Fig 6: Characterization of saphenous nerve innervating the skin.** a) Percent of slowly adapting and rapidly adapting Aβ fibres. b) Representative images of fast conducting NF200 (yellow) positive nerve fibres innervating hair follicles marked by S100b (magenta) and quantification of hair follicles and innervated hair follicles. Scale bar = 50 μm. Each dot represents a mouse, at least 3 images were analyzed per mouse. c) Mean spike rates of slowly adapting mechanosensitive Aβ-fibres in the dynamic phase (in response to ramps of increasing speed) d) (left) Mean spike rates of Aδ-mechano-nociceptive fibres in the static phase in response to increasing force and (right) Mean spike rates of D-hair fibres in the dynamic phase (in response to ramps of increasing speed). Error bars = SEM.

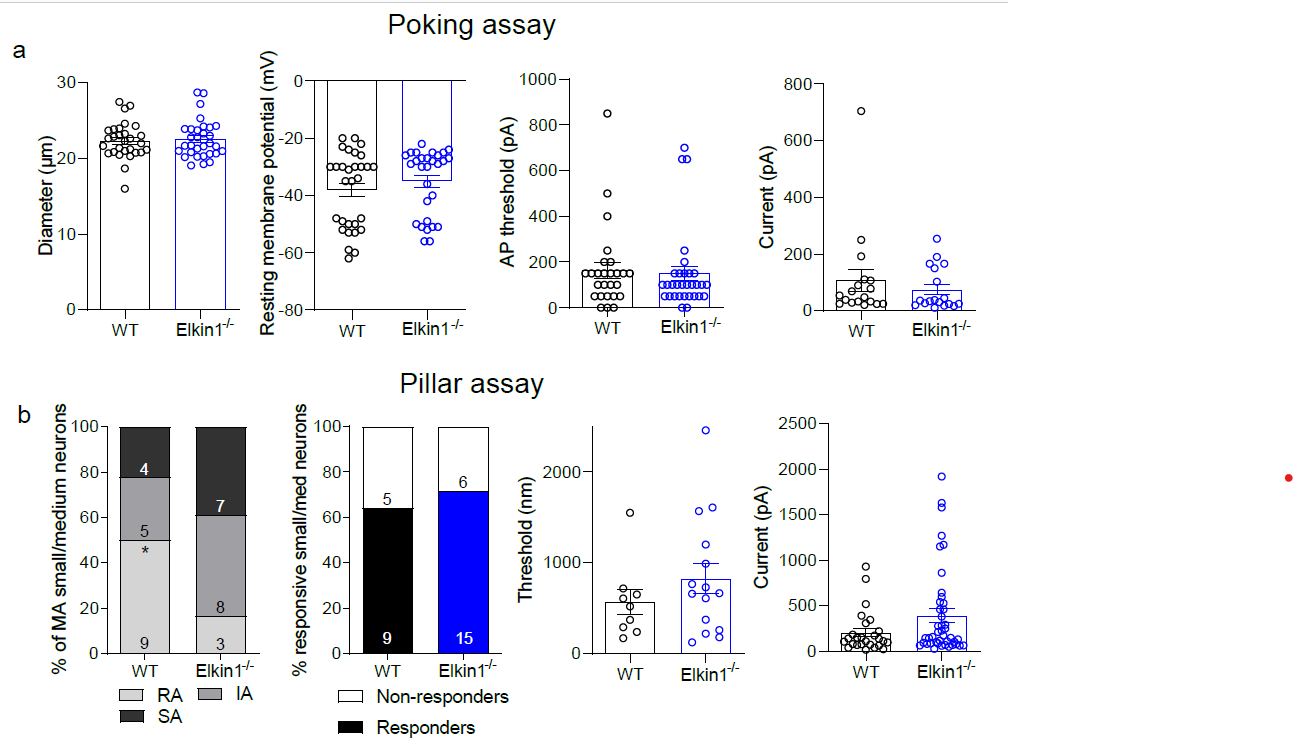

**Extended Data Fig 7: Electrophysiological properties of small/medium sensory neurons.** a) Properties of neurons assessed using poking assay including diameter, resting membrane potential, action potential firing threshold and amplitude of MA currents. Each dot represents a neuron. b) Properties of small/medium neurons as assessed through pillar array assay including percent of MA neurons belonging to rapidly (RA), intermediately (IA) and slowly adapting (SA) category, threshold of MA current activation and max current amplitude of MA currents. Each dot represents a neuron for threshold and each stimulus for current. Data obtained from both male and female mice. Proportions were compared using chi-sq tests. * indicates p < 0.05. Error bars = SEM.

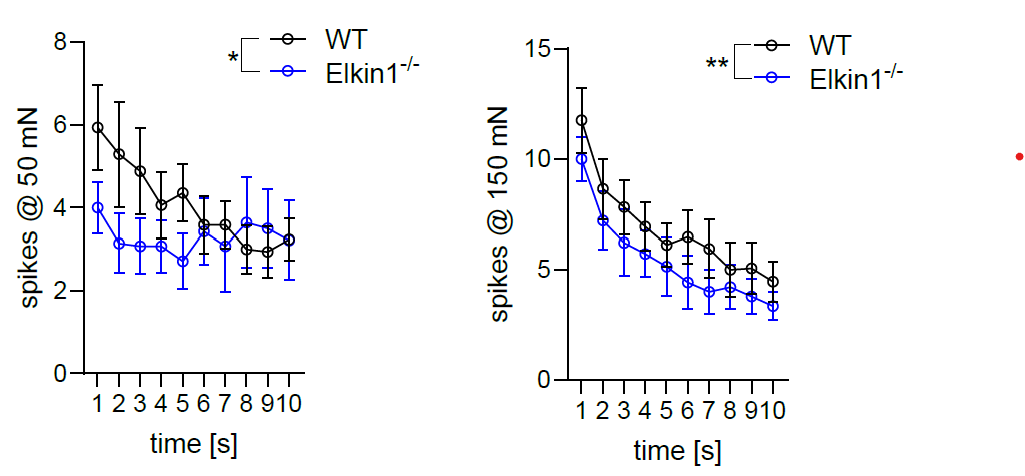

**Extended Data Fig 8: Electrophysiological properties of mechanosensitive C fibres in the saphenous nerve.** Graphs showing absolute number of spikes in response to 50 mN and 150 mN force applied for 10 s. Each dot represents average of n = 17 WT and n = 14 Elkin1-/- fibres from at least 5 mice. Data obtained from both male and female mice. Group comparisons were made using 2 way ANOVA followed by multiple comparison post hoc tests. * indicates p < 0.05, ** indicates p < 0.01. Error bars = SEM.
